# *CellFluxV2* : An Image Generative Foundation Model for Virtual Cell Modeling

**DOI:** 10.64898/2026.01.19.696785

**Authors:** Yuhui Zhang, Yuchang Su, Zoe Wefers, Shiye Su, He Li, Tianhong Li, Chenyu Wang, James Burgess, Alejandro Lozano, Linqi Zhou, Daisy Ding, Jeffrey Nirschl, Emma Lundberg, Serena Yeung-Levy

## Abstract

Building a virtual cell that simulates cellular behavior in silico is a central goal of computational biology. We introduce *CellFluxV2*, an image-generative model that predicts how cell morphologies change in response to chemical and genetic perturbations. A core innovation of *CellFluxV2* is to learn distribution-level transformations from unperturbed to perturbed cells within the same experimental batch using flow matching, enabling it to disentangle true perturbation effects from confounding batch effects. Incorporating three methodological advances, *CellFluxV2* achieves up to a 77% improvement in image fidelity over diffusion- and GAN-based baselines, while maintaining biological fidelity comparable to ground-truth images. Scaling up *CellFluxV2*, we establish the first scaling laws in image-based virtual cell modeling, demonstrating that performance improves consistently with both dataset size and model capacity. Furthermore, the scaled-up model generalizes well to out-of-distribution perturbations and exhibits two novel capabilities: batch-effect correction and cell-state interpolation. Together, these results position *CellFluxV2* as a powerful foundation model advancing the vision of a virtual cell, unlocking novel opportunities for *in silico* drug screening.

---

High-throughput microscopy has become an indispensable tool in drug discovery, enabling researchers to rapidly image cells under thousands of chemical [1–3] and genetic perturbations [4, 5]. Profiling cellular morphological phenotypes across many perturbed conditions can illuminate gene function [6], identify phenotypic signatures of disease [7, 8], predict assay activity [9], and determine the mechanisms of action compounds [10]. In these assays, cells are typically stained with multiple fluorescent dyes or antibodies that label specific molecules, organelles, and cellular components [11–13]. One widely used image-based assay is Cell Painting [14], which uses eight fluorescent dyes to label six major cellular compartments, providing a rich and holistic readout of cell morphology. Because Cell Painting is relatively inexpensive and unbiased, it has become the de facto assay for hypothesis-agnostic screening efforts [10, 13, 15, 16], where the goal is to characterize the effects of large numbers of chemical and genetic perturbations in parallel.

Despite being considered relatively fast, inexpensive, and scalable, high-throughput microscopy and Cell Painting assays can still become costly when applied to combinatorial chemical spaces [17], the 20,000 human protein-coding genes [18], and numerous different cell contexts [19, 20]. As a result, there is growing interest in the development of “virtual cell” models capable of predicting cellular responses to perturbations *in-silico* [21]. Such approaches could facilitate experimental design and accelerate hypothesis generation by reducing the number of experiments required, thereby saving time, cost, and manual labor. In addition, traditional high-throughput microscopy experiments are susceptible to batch effects [22], particularly in large-scale screens that require many rounds of imaging. Although standard batch-correction methods for image-based screens exist, they have been shown to fail on large, diverse datasets [23]. This is problematic because batch effects can confound true perturbational signals, complicating the accurate profiling of new genes and compounds. A virtual cell framework could potentially mitigate this limitation, as in silico predictions would not be subject to experimental batch effects.

Several approaches have been proposed to generate images of perturbed cells. An early method, Mol2Image [24], uses flow-based models conditioned on molecular structure to generate cellular images directly from noise. More recent work has shifted toward conditioning on images of control cells, more explicitly modeling perturbations as transformations of a baseline cellular state. For example, PhenDiff [25] employs a conditional diffusion model to translate cells between conditions, while IMPA [26] uses a style-transfer autoencoder to modify control cells by adding perturbation- and batch-specific “styles.”

However, most existing methods overlook a fundamental experimental constraint: staining and imaging require fixation, which is destructive to cells. Consequently, large-scale perturbation datasets do not contain paired images of the same cell before and after perturbation. To address this mismatch between modeling assumptions and experimental reality, Zhang et al. introduced *CellFlux* [27], which leverages flow matching, a generative modeling framework specifically designed for distribution-to-distribution learning problems. *CellFlux* learns to map the distribution of unperturbed cells to that of perturbed cells within the same experimental batch. This formulation explicitly reflects the biological reality of unpaired data and, when combined with flow matching as a principled solution, enables effective disentanglement of true perturbational effects from confounding biological batch effects.

In this work, we present *CellFluxV2*, an extension of the original framework. We show that incorporating three additional techniques, latent-space modeling, two-stage training, and noisy interpolants [28], efficiently mitigates the impact of data sparsity. As a result, *CellFluxV2* achieves improved image quality and biological fidelity, significantly outperforming CellFlux, IMPA, and other baseline models. Building on these gains, we scale *CellFluxV2* by training a larger model on more data, resulting in additional performance improvements in image quality as well as enhanced out-of-distribution generalization to unseen batches and compounds. These results establish the first empirical scaling laws for biological perturbation modeling, mirroring trends previously observed in large language models [29]. Additionally, we demonstrate that *CellFluxV2* can faithfully recapitulate batch effects in its generated perturbed images. By matching the batch characteristics of generated images to those of the comparison set, *CellFluxV2* effectively isolates true perturbational signals, leading to improved mechanism-of-action prediction. Finally, we show that *CellFluxV2* learns smooth, bidirectional transitions between control and perturbed states and that the transitional states match the true morphology of perturbed cells at intermediate time points. These findings highlight an opportunity to use flow-matching approaches for cell-state interpolation and suggest a new avenue for modeling morphological trajectories over time.

Taken together, these results establish *CellFluxV2* as a generalizable and scalable approach for modeling cellular perturbations from unpaired image data within a principled distribution-to-distribution framework. This work lays the foundation for the next generation of image-based *in-silico* perturbation models and represents a substantive step toward the development of a virtual cell.

## Results

### Distribution-wise Transformation using Flow Matching

*CellFluxV2* is an image-based generative model that predicts how cellular morphology changes in response to chemical or genetic perturbations in silico. By modeling these perturbation-induced transitions, *CellFluxV2* has the potential to accelerate phenotypic screening and enable downstream applications in basic biology, drug discovery, and personalized medicine (Fig. 1a).

**Fig. 1.**
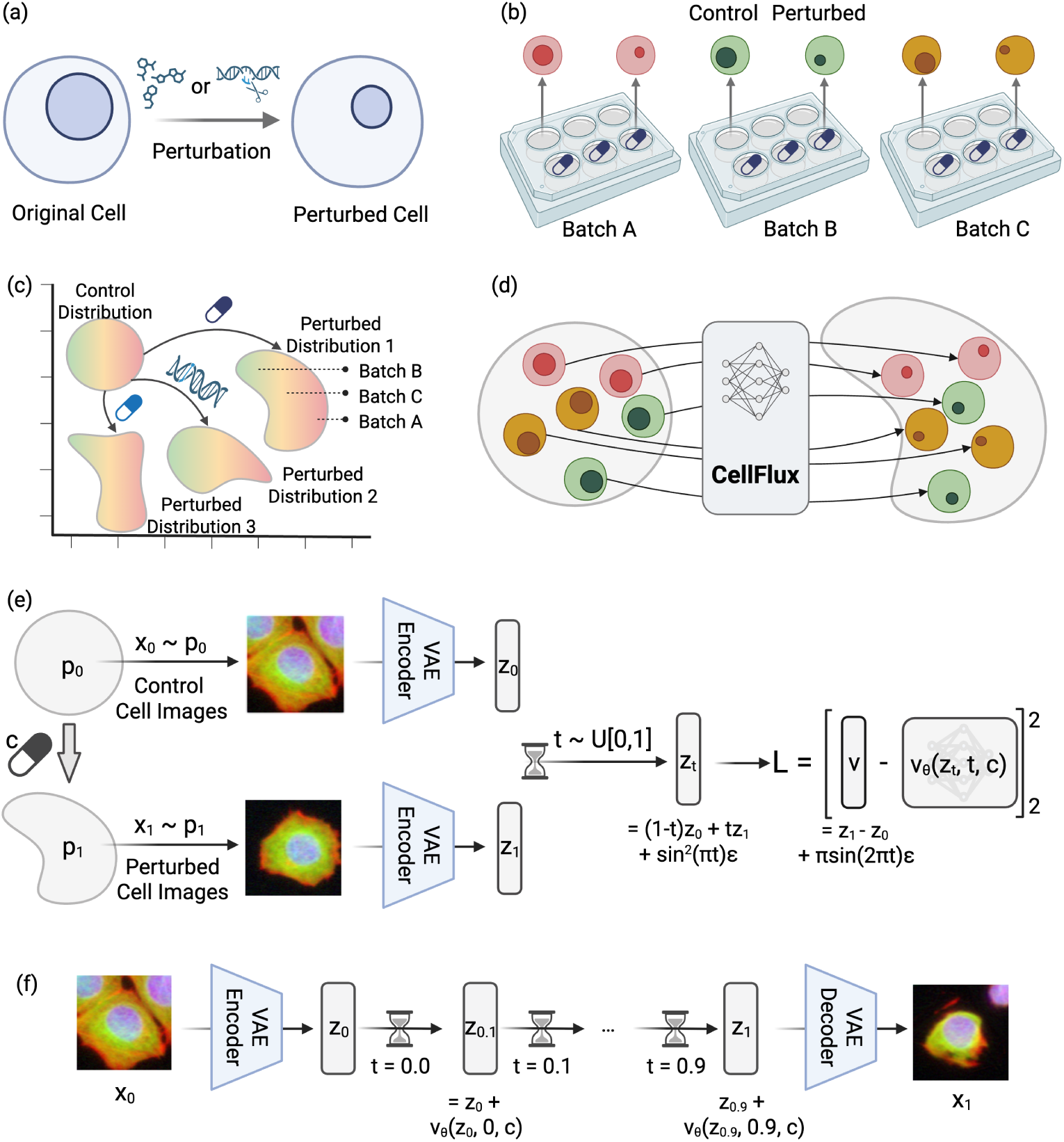
Overview of *CellFluxV2*. *(a) Objective. CellFluxV2* aims to predict changes in cell morphology induced by chemical or gene perturbations *in silico*. In this example, the perturbation effect reduces the nuclear size. *(b) Data.* The dataset includes images from high-content screening experiments, where chemical or genetic perturbations are applied to target wells, alongside control wells without perturbations. Control wells provide prior information to contrast with target images, enabling the identification of true perturbation effects (e.g., reduced nucleus size) while calibrating non-perturbation artifacts such as batch effects—systematic biases unrelated to the perturbation (e.g., variations in color intensity). *(c) Problem formulation. CellFluxV2* formulates the task as a distribution-to-distribution problem (many-to-many mapping), where the source distribution consists of control images, and the target distribution contains perturbed images within the same batch. *(d) Flow matching. CellFluxV2* leverages flow matching, a state-of-the-art generative framework for distribution-to-distribution transformation. The model learns a neural network that approximates a velocity field, enabling continuous transformation of the source distribution into the target by solving an ordinary differential equation (ODE). Compared to *CellFluxV1*, we introduce three key improvements: latent space modeling, two-stage training, and noisy interpolants. Together, these techniques effectively mitigate the data sparsity issue in distribution learning. *(e) CellFluxV2 algorithm.* During training, the neural network *v_θ_* learns a velocity field mapping control cell images (*x*_0_ ∼ *p*_0_) to perturbed images (*x*_1_∼ *p*_1_) in latent space. Intermediate states *x_t_* are sampled along the linear interpolation between *x*_0_ and *x*_1_ with *t* ∼ *U* [0, 1] and noisy augmentation. The loss *L* matches the predicted velocity *v_θ_*(*x_t_, t, c*) to the true displacement (*x*_1_ − *x*_0_). The method is trained in two stages: the first stage goes from noise to the target, and the second stage goes from the control to the target. During inference, the trained field *v_θ_* transforms a control state *x*_0_ toward the perturbed state by numerically integrating the ODE over time steps *t* = 0, 0.1*, …,* 1, updating the state at each step using the learned velocity.

A key design principle of *CellFluxV2* is that it learns *distribution-to-distribution transformations*: rather than generating perturbed cells de novo, the model learns to transform the distribution of unperturbed control cells into that of perturbed cells. This choice is motivated by two fundamental properties of high-throughput imaging data.

#### Inappropriate noise-to-distribution formulation due to batch effect

First, batch effects make a noise-to-distribution formulation inappropriate. High-content imaging experiments are typically performed across multiple plates, each representing a separate experimental batch. Cell images from different batches can differ substantially in color intensity, contrast, cell density, and other acquisition-specific factors. If a model were to generate perturbed cells directly from Gaussian noise, it would be unable to disentangle true biological effects from these batch-specific artifacts. As a result, the model might incorrectly attribute differences due to batch variability—for example, changes in illumination or staining intensity—to the perturbation itself, rather than capturing the true morphological changes such as nuclear shrinkage (Fig. 1b). Conditioning on an observed unperturbed cell state is therefore essential for calibrating away batch effects. Prior GAN- and diffusion-based approaches often cannot exploit this information because they generate images solely from noise and cannot start from an observed unperturbed cell.

#### Impossible one-to-one formulation due to destructive nature of fixed imaging

Second, the destructive nature of fixation and staining precludes a one-to-one formulation. Although we require the basal state of a cell to correct batch effects, it is impossible to obtain the exact pre-perturbation image of any given cell: cell painting requires fixation, which kills the cells and prevents the same cell from subsequently experiencing a perturbation. Experimental workflows instead rely on control cells—unperturbed cells obtained from the same batch—but there is no one-to-one correspondence between individual control cells and perturbed cells. Consequently, the task must be understood as a distribution-to-distribution transformation, where the model learns how the overall population of control cells differs from the perturbed population, thereby averaging out batch effects and isolating the true phenotypic signal.

Together, these considerations make a distribution-to-distribution formulation not only natural but necessary (Fig. 1c).

#### Flow matching for distribution-to-distribution

Recent advances in generative modeling techniques, such as flow matching, offer a principled solution for the distribution-to-distribution problem. Flow matching extends the intuition behind diffusion models by learning a continuous velocity field that transports one empirical distribution into another, without requiring either distribution to follow a Gaussian or simplified structure. Although flow matching has achieved state-of-the-art generation performance across domains such as images, videos, and biological sequences, most mainstream applications use it in restricted forms—either noise-to-distribution (as in diffusion models) or one-to-one image editing—where its full flexibility is not needed. As a result, its capacity to learn rich distribution-to-distribution mappings has remained largely untapped. This gap makes it a natural and powerful fit for our setting (Fig. 1d). The training and inference procedures for the flow-matching algorithm are provided in Fig. 1e and Fig. 1f, with more details in the Methods section.

*CellFluxV2* leverages the full distribution-to-distribution formulation of flow matching to model perturbation effects more faithfully. We demonstrate in later sections that this design substantially outperforms GAN and diffusion baselines (Fig. 2). Beyond advancing virtual cell modeling, *CellFluxV2* also offers one of the first real-world demonstrations of flow matching’s ability to perform general distribution transformations without assuming a Gaussian source distribution, underscoring its broader utility for machine-learning research.

**Fig. 2.**
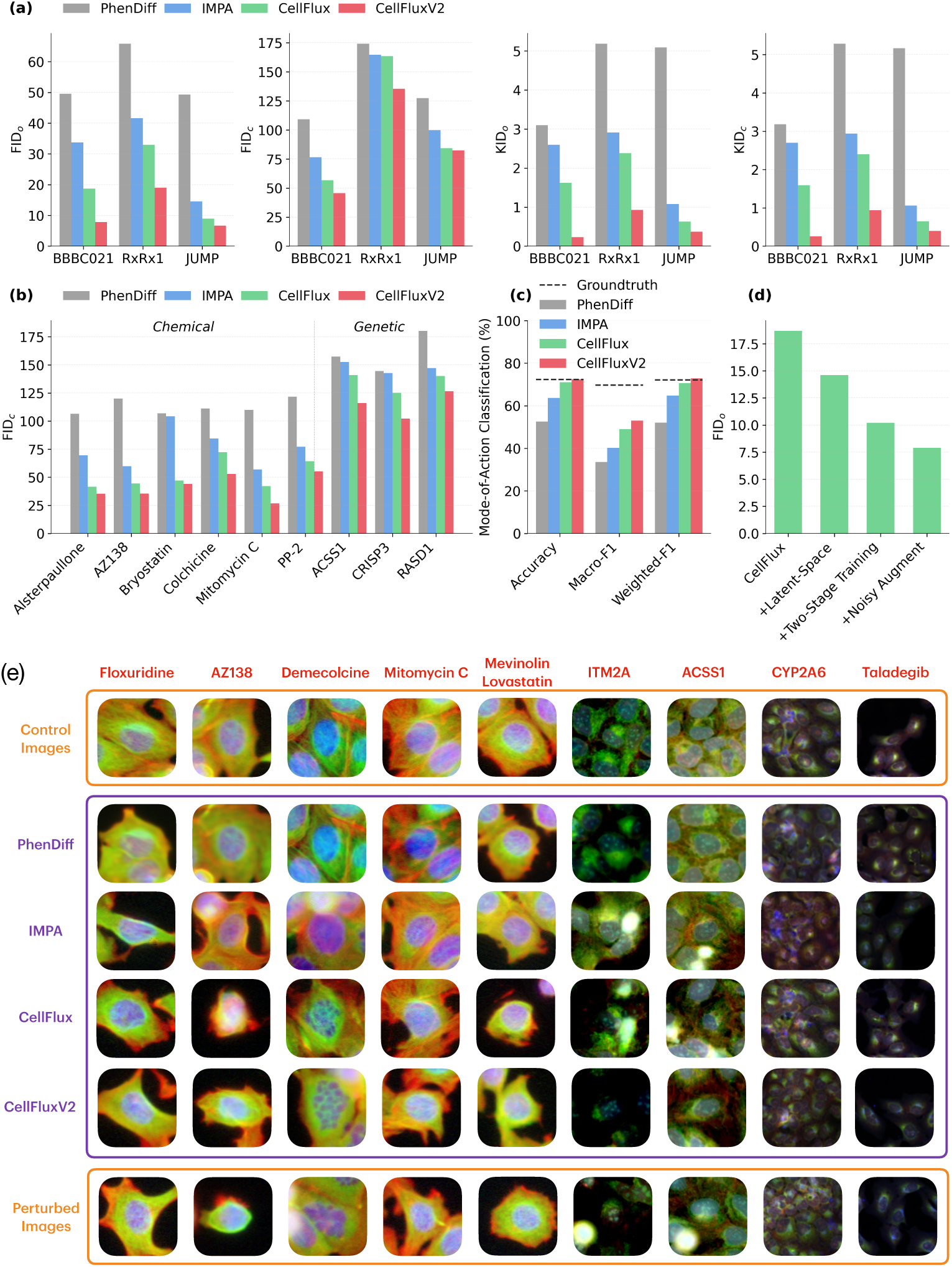
*CellFluxV2* outperforms baselines by a large margin. *(a) Image fidelity. CellFluxV2* achieves superior image fidelity, evidenced by significantly lower overall and conditional FID/KID. *(b) Per perturbation results.* For six representative chemical perturbations and three genetic perturbations, *CellFluxV2* generates significantly more accurate images that better capture the perturbation effects than other methods, as measured by the FID score. *(c) Biological fidelity. CellFluxV2* achieves superior biological fidelity, evidenced by higher and near–ground-truth mode-of-action (MoA) prediction accuracy. *(d) Method ablation.* The superior performance of *CellFluxV2* arises from innovations that substantially improve flow matching for distribution-to-distribution transformations, including latent-space modeling, two-stage training, and noisy interpolants. *(e) Qualitative results.* Qualitative analyses further confirm the superior visual and biological fidelity of *CellFluxV2*.

### State-of-the-art Performance

*CellFluxV2* substantially outperforms prior methods across three perturbation datasets—BBBC021 (chemical), RxRx1 (genetic), and JUMP (combined)—demonstrating markedly improved visual fidelity and biological accuracy.

#### CellFluxV2 achieves superior image fidelity, evidenced by significantly lower overall and conditional FID/KID

We evaluated image quality using the Fréchet Inception Distance (FID) and Kernel Inception Distance (KID), which measure discrepancies between generated and real images (lower values indicate higher fidelity). We report each metric in two settings: (1) *overall* quality (denoted FID*_o_* / KID*_o_*), comparing all generated images to all real perturbed images, and (2) *conditional* quality (denoted FID*_c_* / KID*_c_*), comparing generated and real perturbed images for each perturbation type and averaging across perturbations.

*CellFluxV2* achieves overall FID*_o_* scores of 7.9, 19.0, and 6.7 on BBBC021, RxRx1, and JUMP, respectively—representing improvements of up to 76% over the previous best method, *CellFluxV1* (18.7, 33.0, and 9.0), and similarly outperforming diffusion- and GAN-based baseline methods (Fig. 2a). Conditional FID*_c_* results further confirm that *CellFluxV2* captures fine-grained, perturbation-specific morphological patterns rather than generating generic cell-like textures. Detailed conditional FID results for representative perturbations are provided in Fig. 2b. The same performance trends hold for KID*_o_* and KID*_c_* (Fig. 2a), underscoring the robustness of the gains in visual fidelity.

#### CellFluxV2 achieves superior biological fidelity, evidenced by higher and near-real-image MoA prediction accuracy

To quantify biological fidelity, we assessed whether the generated images preserve morphological features relevant to the mode of action (MoA) of each chemical perturbation, where MoA refers to the specific biological pathway or molecular mechanism through which a compound exerts its effect. Because the MoA classifier (a neural network classifier trained on real images) is imperfect—achieving 72.4% accuracy and 72.1% weighted F1 on real BBBC021 images—performance close to this empirical ceiling indicates strong biological realism. *CellFluxV2* achieves 72.5% accuracy and 72.9% weighted F1, effectively matching the classifier’s performance on real images, with only a minor drop in macro F1. These results represent improvements of 8.8 percentage points in accuracy (72.5% vs. 63.7%) and 8.1 points in weighted F1 (72.9% vs. 64.8%) over the diffusion- and GAN-based baselines. Together, these findings demonstrate that *CellFluxV2* preserves the key morphological signatures required to distinguish perturbation mechanisms (Fig. 2c).

#### Visual inspections further confirm the superior visual and biological fidelity of CellFluxV2

Fig. 2e presents visual comparisons between *CellFluxV2*, *CellFluxV1*, and strong GAN- and diffusion-based baselines (IMPA and PhenDiff). *CellFluxV2* consistently generates sharper cellular structures, more realistic subcellular organization, and clearer perturbation-specific phenotypes. For example, *AZ138* is an Eg5 inhibitor, which induces *cell death and shrinkage*, *CellFluxV2* accurately reproduces *cell shrinkage*. Likewise, for perturbation *Demecolcine*, which destabilizes microtubules, resulting in *smaller, fragmented nuclei*, *CellFluxV2* captures *nuclei change*. These qualitative results reinforce the quantitative gains in both visual fidelity and biological accuracy.

#### The superior performance of CellFluxV2 arises from innovations that substantially improve flow matching for distribution-to-distribution transformations

As mentioned earlier, although flow matching is designed for arbitrary distribution transformations, most current use cases assume a Gaussian source distribution, similar to diffusion models. In our work, when applying flow matching to true distribution-to-distribution transformation, we encounter a central challenge: data sparsity. To mitigate this sparsity, we introduce three innovations that work together to improve performance: (1) latent space modeling, (2) two-stage training, and (3) noisy interpolants. A brief overview is provided below, with full technical details in the Methods section.

- *Latent-space modeling:* We train a variational autoencoder (VAE) to embed high-dimensional pixel space into a compact, semantically meaningful latent space. Training the velocity field in this lower-dimensional manifold reduces data sparsity, improves generalization, decreases FID by 22%, and significantly accelerates training (Fig. 2d).
- *Two-stage training:* Because our data overall contains far fewer control cells than perturbed cells, the source distribution is especially sparse. To mitigate this, we stabilize optimization via a warm-up phase in which the model first learns to map Gaussian noise to the target distribution of perturbed cells. After this initialization, we fine-tune the model to transform control-cell distributions into their perturbed counterparts. This two-stage approach mitigates overfitting and yields a further 30% improvement in FID (Fig. 2d).
- *Noisy interpolants:* To combat sparsity in high-dimensional spaces, we inject Gaussian noise into rectified path interpolants, effectively enriching the training trajectories and providing additional support in regions of the latent space poorly covered by image samples. This augmentation improves robustness and contributes to the final 25% performance gains (Fig. 2d).

### Scaling laws

Scaling laws, defined as systematic improvements in performance as model and dataset size increase, have been central to the progress of modern AI systems such as GPT-style large language models. For virtual cell modeling, establishing whether similar principles apply is important because it would clarify how far performance can be pushed with larger architectures and richer microscopy datasets, and whether continued investment in scale is scientifically justified. Motivated by this, we used the state-of-the-art *CellFluxV2* architecture to examine whether analogous behaviors emerge in this domain. By jointly scaling model capacity and the amount of training data, we observed consistent gains across both axes, indicating that comparable scaling laws indeed hold for virtual cell modeling.

#### Scaling with data improves performance

To study data scaling effects, we processed the full JUMP-pilot dataset, which comprises 11.6 million images spanning 817 perturbations and exceeds 1 terabyte in size. We partitioned the dataset into four subsets based on in-distribution or out-of-distribution perturbations and control images (details provided in the Appendix). Models were trained on five randomly sampled data fractions—3%, 10%, 30%, 50%, and 100%—each subset containing all perturbations.

On the in-distribution evaluation set, models achieved FID_o_ scores (computed on 5000 sampled images) of 18.3, 9.6, 7.1, 6.7, and 6.0, respectively, corresponding to a 67% improvement (Fig. 3a). These results highlight the value of high-throughput imaging platforms, which can generate terabyte- to petabyte-scale datasets that substantially enhance virtual cell modeling.

**Fig. 3.**
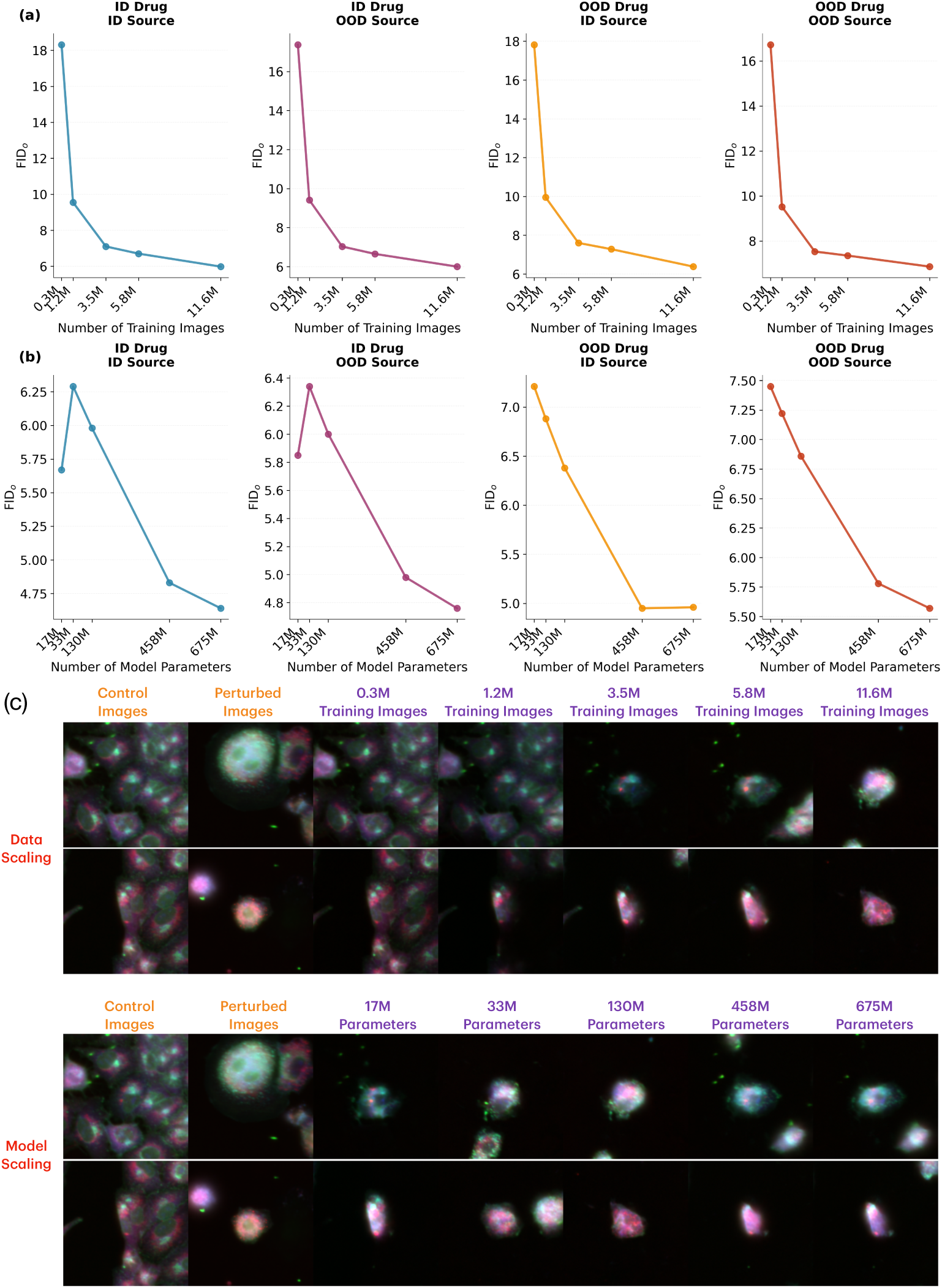
Scaling up *CellFluxV2* reveals the first scaling laws in virtual cell modeling. *(a) Data scaling.* Scaling with data (from 3% to 100% using base model) improves performance. *(b) Model scaling.* Scaling with model size (from tiny to extra large using 100% data) improves performance. Together, we establish the first empirical scaling laws in virtual cell modeling, demonstrating that model performance improves substantially with both increasing data scale and model size. Beyond in-distribution performance, scaling also enhances the model’s ability to generalize to out-of-distribution image sources and perturbations, highlighting it spotential as a robust foundation for virtual cell modeling. *(c) Qualitative results.* Qualitative analyses (using colchicine as the perturbation) further confirm improved generation quality with increased data and model scaling (colchicine causes cells to adopt more spherical or rounded shapes).

#### Scaling with model size improves performance

To investigate model-capacity scaling, we used a Transformer backbone to learn the velocity field in flow matching, motivated by the strong scalability of Transformer architectures in general-purpose image generation. We trained five models of increasing size—tiny (17M parameters), small (33M), base (130M), large (458M), and extra-large (675M)—all using the full dataset.

On in-distribution evaluations, the models achieved FID_o_ scores (computed on 5000 sampled images) of 5.7, 6.3, 6.0, 4.8 and 4.6, demonstrating a 27% improvement (Fig. 3b). Although the tiny model performs slightly better than the small and base models in the in-distribution setting, this comes at the cost of significantly worse performance in out-of-distribution settings. These findings align with trends in large-scale AI, where performance continues to increase with model capacity, particularly when sufficient data are available.

#### Visual inspections further confirm the scaling effect

Fig. 3c presents visual comparisons of generated images across different dataset sizes and model scales under colchicine as the perturbation. Colchicine alters cell morphology by binding to tubulin and inhibiting microtubule polymerization, which destabilizes the cell’s cytoskeletal framework. This disruption causes the cell to lose its specific structural integrity and retract into a spherical or rounded shape. We observe that the generated images become more similar to the ground-truth perturbed images (spherical or rounded shape) as both the training data and model size increase. These qualitative results reinforce the quantitative gains in visual fidelity and biological accuracy, highlighting the importance of scaling laws.

#### Scaling laws in virtual cell modeling and their implications

Together, these experiments reveal the first scaling laws in virtual cell modeling: performance improves monotonically with both larger datasets and larger model architectures. This mirrors the pattern observed in language and vision models, where scaling laws have guided the development of increasingly capable systems. The emergence of similar behavior in virtual cell modeling has broad implications: coupled with rapid advances in generative modeling and high-throughput biotechnology, these scaling trends bring us significantly closer to the long-standing goal of achieving a comprehensive “virtual cell.”

### Out-of-distribution Generalization

As *CellFluxV2* achieves strong performance in in-distribution settings, we next evaluate whether it can generalize to unseen, out-of-distribution (OOD) scenarios—a core requirement for a practical virtual cell model. A virtual cell is substantially more useful if it can operate reliably under conditions not encountered during training, such as previously unseen drugs, genetic perturbations, or imaging environments from different laboratories. Such capabilities are essential for reducing wet-lab experimentation costs and enabling broad utility in biomedical research and drug discovery. We evaluated *CellFluxV2* on OOD perturbations, OOD imaging sources, and their combination (Fig. 4) and found that it generalizes remarkably well. In the most challenging setting—simultaneous OOD perturbations and OOD imaging conditions—the model’s FID remains comparable to its in-distribution performance. Notably, scaling both model capacity and training data consistently improves generalization.

**Fig. 4.**
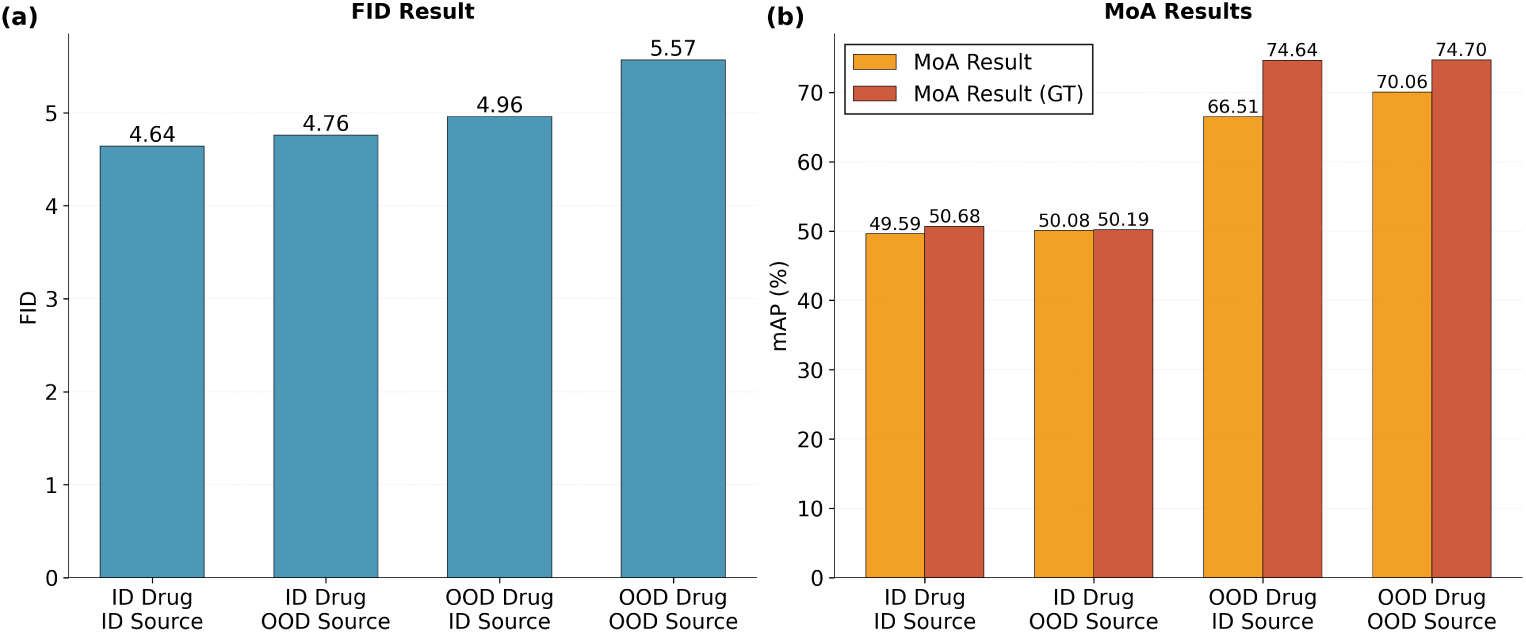
*CellFluxV2* demonstrates robust generalization to out-of-distribution (OOD) perturbations and imaging conditions. *(a) Image fidelity metrics.* The model generalizes to OOD imaging conditions and perturbations, achieving image fidelity metrics (FID / KID) comparable to in-distribution performance. *(b) Biological fidelity metrics.* The model generalizes to OOD imaging conditions and perturbations, achieving biological fidelity metrics such as mode-of-action (MoA) mAP similar to in-distribution results and close to the ground truth.

#### Model generalizes to OOD plate batches

We first assess the scenario in which the perturbation type (denoted as P) is observed during training, but the control images (denoted as I) originate from an unseen source (e.g., a plate never used in training), resulting in OOD variation due to batch effects. As shown in Fig. 4a/b, *CellFluxV2*-XL achieves 4.8 FID*_o_* and 0.5008 MoA mAP (real image MoA mAP is 0.5019) in this *OOD^I^ –ID^P^* setting (see the Methods section for MoA mAP calculation). Importantly, we again observe clear scaling trends: increasing both model size and training data yields monotonic improvements on this OOD benchmark. These results demonstrate that *CellFluxV2* robustly generalizes to new imaging conditions, enabling reliable predictions on images acquired from other laboratories or experimental setups.

#### Model generalizes to OOD perturbations

Next, we evaluate the complementary scenario in which control images are from the training distribution, but the perturbations are entirely new—for example, unseen drugs or gene edits. As shown in Fig. 4a/b, *CellFluxV2*-XL attains 5.0 FID*_o_* and 0.6651 MoA mAP (real image MoA mAP is 0.7464) in this *ID^I^ –OOD^P^* setting. Similar scaling laws emerge, with performance improving as model size and dataset size increase. These findings indicate that *CellFluxV2* can generalize to novel perturbations, enabling prediction for new compounds or genetic interventions and thereby reducing the need for costly wet-lab experiments.

#### Model generalizes to both OOD images and perturbations

Finally, we consider the most challenging regime, in which both the perturbation and the imaging conditions are unseen during training. As shown in Fig. 4a/b, *CellFluxV2*- XL achieves 5.6 FID*_o_* and 0.7006 MoA mAP (real image MoA mAP is 0.7470) in this *OOD^I^ –OOD^P^* setting. We again observe consistent scaling behavior. Together, these results show that *CellFluxV2* is well-prepared for extreme OOD scenarios, bringing us closer to a truly generalizable virtual cell model.

### New Capability: Batch Effect Correction

As discussed, a central contribution of *CellFluxV2* is its formulation of cell-perturbation modeling as a *distribution-to-distribution transformation* problem. This allows the model to explicitly account for the well-known *biological batch effect*, in which images from different experimental batches can differ substantially in intensity, morphology, or cell density. In the following sections, we demonstrate this capability using (i) qualitative visualizations showing that *CellFluxV2* preserves batch-specific characteristics when conditioned on batch inputs, (ii) PCA analyses demonstrating that generated images cluster with their corresponding batch, and (iii) quantitative mode-of-action (MoA) metrics showing that conditioning on the correct batch substantially improves predictive accuracy.

#### Qualitative results: Batch-conditioned generation

We applied *CellFluxV2* to generate perturbed cell images conditioned on two batches that differ markedly in overall intensity and cell density (Fig. 5a). The generated images faithfully reproduce the batch-specific appearance—including characteristic intensity levels and density patterns—while still expressing the true perturbation signature (e.g., shrinkage of cell boundaries). In contrast, GAN- or diffusion-based baselines that do not model distribution-to-distribution mappings must generate images from Gaussian noise; as a result, they are unable to control for batch effects, and their generated images often show cell density changes. Without access to the true control image, such artifacts could easily be misinterpreted as biological effects rather than experimental confounders.

**Fig. 5.**
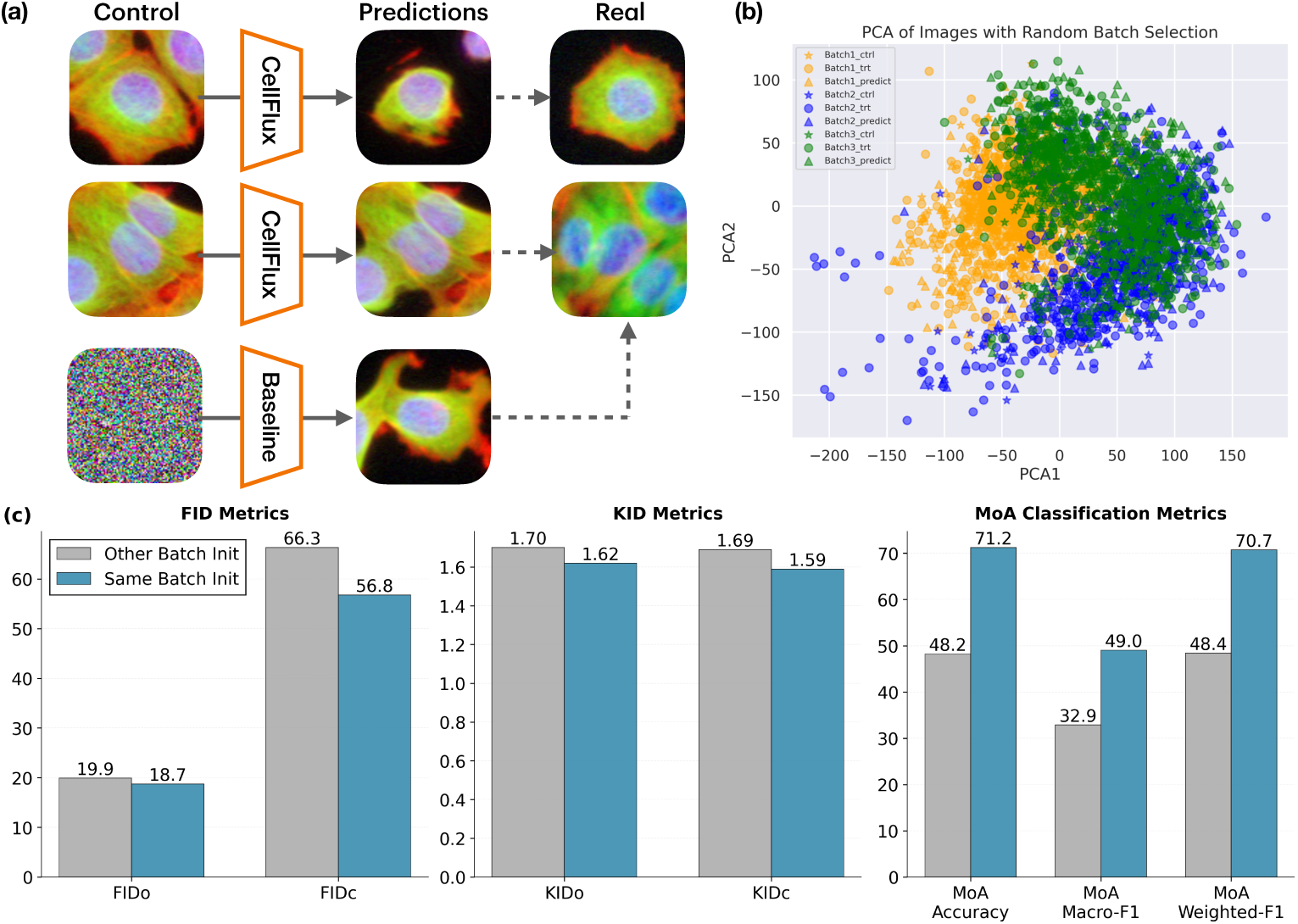
*CellFluxV2* enables the novel capability of batch effect correction. *(a) Qualitative Results from Different Batch. CellFluxV2* initializes with control images, enabling batch-specific predictions. Comparing predictions from different batches highlights actual perturbation effects (smaller cell size) while filtering out spurious batch effects (cell density variations). *(b) PCA Visualization of Cells from Different Batch.* Generated, ground-truth, and control images were projected into a low-dimensional space using PCA, with points colored by batch identity. *CellFluxV2*-generated images cluster closely with their corresponding batches, overlapping with both ground-truth and control cells from the same batch. *(c) MoA Accuracy Results.* When conditioning on the correct batch, *CellFluxV2* generated images yield significantly higher MoA classification accuracy compared to conditioning on different batches.

This ability to preserve batch structure while isolating genuine perturbation effects is a fundamental property of *CellFluxV2*, enabling both substantial performance gains and improved interpretability by providing generated cell images that reflect biological signals instead of batch artifacts.

#### Qualitative results: PCA visualization

To further assess whether *CellFluxV2* captures batch-specific structure, we projected generated images, real perturbed images, and control images into a low-dimensional space using PCA and colored points by batch identity (Fig. 5b). Images generated by *CellFluxV2* cluster tightly with the corresponding batch, overlapping with both the real perturbed and control images from that batch. This confirms that *CellFluxV2* preserves the underlying batch distribution and reproduces batch effects in a controlled and biologically consistent manner.

#### Quantitative results: MoA prediction

We generated perturbed images by conditioning *CellFluxV2* either on the correct batch or on mismatched batches, and evaluated the resulting images using mode-of-action (MoA) classification. Conditioning on the correct batch yields 71.2% MoA accuracy, whereas conditioning on different batches yields only 48.2% accuracy (Fig. 5c). The large performance gap demonstrates that proper handling of batch effects is essential: without batch-aware generation, MoA predictions become dominated by batch artifacts rather than genuine perturbation biology.

### New Capability: Cell State Interpolation

A novel capability introduced by *CellFluxV2* is its ability to interpolate between two cell states using a continuous velocity field learned through a flow-matching algorithm. This enables us to generate smooth trajectories that may reflect biologically meaningful transitions—for example, accurately predicting the intermediate cell states occurring halfway through a perturbation time course.

#### Continuous velocity fields learned through flow matching enable interpretable interpolation trajectories

The flow-matching algorithm learns a continuous velocity field that maps one cell-image distribution to another. As a result, *CellFluxV2* can perform smooth, bidirectional interpolation between two cellular states. Starting from a control image, we iteratively follow the learned velocity field to generate intermediate images along the trajectory. As illustrated in Fig. 6a, these intermediate states evolve in a semantically coherent manner; for perturbations that reduce cell size, the interpolated cells gradually become smaller along the trajectory. Such trajectory visualizations are particularly valuable because they are difficult to obtain experimentally: Cell Painting assays require cell fixation, preventing time-lapse imaging of the same cell. The interpolated trajectories, therefore, provide a unique window into the underlying biological progression and can help researchers infer dynamic cellular processes.

**Fig. 6.**
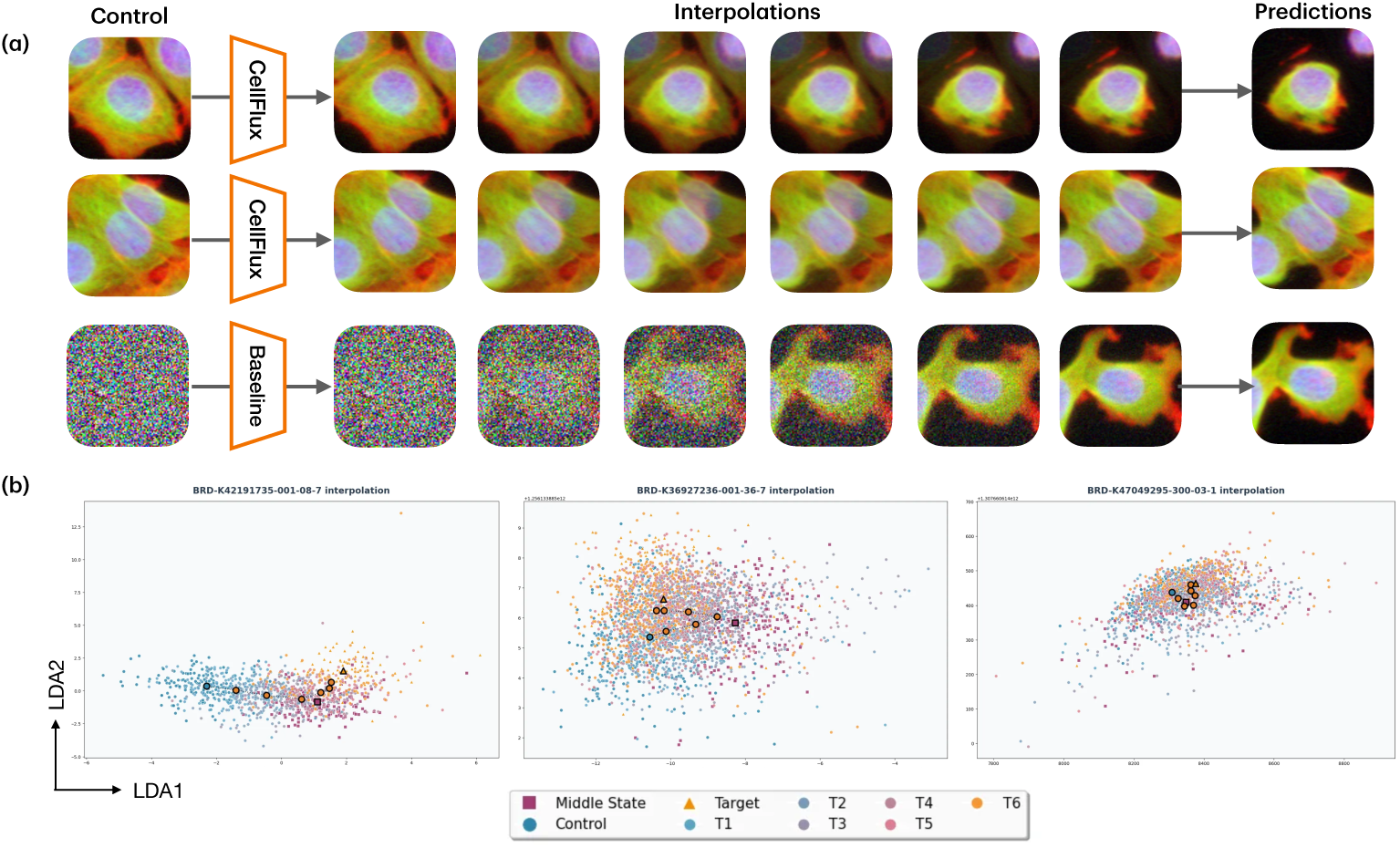
*CellFluxV2* enables the novel capability of cell state interpolation. *(a) Qualitative Results of Interpolation.*Using a continuous velocity field, *CellFluxV2*, perform smooth, bidirectional interpolation between two cell states, which provides a unique window into dynamic biological progression. *(b) PCA Visualization of Interpolated Trajectories.* We projected both the interpolated images and the real images of the CellProfiler feature into a low-dimensional space using PCA. *Control*, *Target*, and *Middle State* represent CellProfiler embedding of control cells, cells subjected to 48-hour and 24-hour perturbation. Other colored points (*T*_1_ − *T*_6_) correspond to generated cells collected at intermediate time points. The interpolated images clearly depict a smooth transition from Control to Target, with intermediate timestep samples clustering closely with the ground-truth Middle State cells. Axes show the first two principal components with explained variance in parentheses.

#### PCA visualization confirms that interpolated trajectories reflect biological progression

To assess whether the interpolated trajectories capture true biological dynamics, we performed a case study interpolating between control cells and cells subjected to a 48-hour perturbation. We also collected real perturbed images at the 24-hour midpoint. We projected both the interpolated images and the real images into a low-dimensional space using PCA (Fig. 6b). The interpolated images at *t* = 0.5 cluster closely with the 24-hour real perturbed cells, and the trajectory progresses smoothly from the control population toward the 48-hour target state. This clear correspondence demonstrates that the *CellFluxV2* interpolation trajectory is biologically meaningful, suggesting a flow-matching as a novel approach for studying cell-state transitions and dynamics.

## Discussion

High-throughput microscopy provides rich and scalable readouts of cellular responses to genetic and chemical perturbations, but its experimental cost and complexity motivate the development of predictive *in-silico* alternatives. In this work, we introduce *CellFluxV2*, a flow-matching–based generative model that synthesizes realistic images of chemically and genetically perturbed cells. *CellFluxV2* formulates perturbation modeling as a distribution-to-distribution transformation problem and applies flow matching as a principled solution for learning from unpaired data. To address data sparsity, *CellFluxV2* incorporates multiple training strategies that improve image fidelity and enhance downstream biological performance, including mechanism-of-action prediction. We show that *CellFluxV2* outperforms prior methods in both image quality and biological accuracy, and that scaling model size and dataset size leads to further improvements in image realism and out-of-distribution generalization.

Beyond methodological advances, we demonstrate the practical utility of *CellFluxV2* across several challenging settings. The model accurately generates perturbed cellular phenotypes for unseen experimental batches and for a subset of previously unseen compounds. By adopting a training scheme that transforms control cells into perturbed cells within the same experimental batch, *CellFluxV2* learns to disentangle true perturbational effects from batch-specific artifacts. We show that this property can be leveraged to mitigate batch effects by generating perturbed images within a common batch, resulting in improved mechanism-of-action prediction. In addition, *CellFluxV2* learns a continuous velocity field over cellular states, enabling smooth and biologically meaningful interpolation between control and perturbed conditions.

Despite these advances, several limitations remain. Although *CellFluxV2* is trained on a comparatively large dataset, it is exposed to only 306 compounds and 161 genes, representing a small fraction of chemical space and druggable targets. Consequently, the model is unlikely to accurately predict perturbations that induce morphological phenotypes not observed during training. Moreover, while our treatment of batch effects captures plate-to-plate variability, real-world datasets often include more severe sources of variation, such as differences across protocols, microscopes, and laboratories. Finally, our experiments focus on two cell lines from the JUMP pilot dataset; however, phenotypic responses to perturbations can vary substantially across cell types, limiting direct generalization to new cellular contexts.

Looking ahead, *CellFluxV2* could be further scaled to larger and more chemically diverse datasets, such as JUMP-CP [13], EU-OPENSCREEN [10], or RxRx3 [16]. Given the scaling trends observed in this work, we expect performance to continue improving with increased data and model capacity. Training on broader chemical spaces would further alleviate data sparsity and enhance generalization to unseen perturbations. In addition, larger datasets spanning multiple laboratories, microscopes, and experimental conditions would provide an opportunity to more rigorously evaluate and extend *CellFluxV2*’s ability to model and mitigate complex batch effects. Future work could also explore transfer across cell types, for example, by training on one cell line and adapting the model to another.

In addition, *CellFluxV2* currently relies on relatively simple perturbation encodings, using Morgan fingerprints for small molecules and gene2vec embeddings for genetic perturbations. More expressive representations, such as encodings of 2D molecular structures and embeddings from protein or DNA language models, could enable richer modeling of perturbation mechanisms and improve generalization to unseen compounds and genes. Further, the ability of *CellFluxV2* to perform smooth cell-state interpolation offers an opportunity to move beyond static perturbation endpoints and toward modeling continuous morphological trajectories. Extending flow-matching approaches to time-lapse imaging data could enable *in-silico* exploration of temporal cellular dynamics. Finally, extending *CellFluxV2* to a multimodal setting could enable joint modeling of imaging and transcriptomic perturbation data, leveraging the growing number of large-scale transcriptomic screens alongside microscopy datasets. Such a model could capture perturbational effects that manifest in only one modality and, if successful, generate synthetic paired cellular images and gene expression profiles, which are not currently achievable experimentally.

Overall, *CellFluxV2* positions flow matching as a robust and flexible foundation for image-based modeling of cellular perturbations. Continued advances in scale, biological representation, and data diversity are likely to further enhance performance and practical utility, bringing the field closer to the goal of accurate and generalizable virtual cell models.

## Methods

### Problem Formulation

We first introduce the objective, data, and mathematical formulation of cellular morphology prediction.

#### Objective

Let X denote the cell image space and C the perturbation space. Let *p*_0_ represent the original cell distribution and *p*_1_ represent the distribution of cells after a perturbation *c* ∈ C. Cellular morphology prediction aims to learn a generative model *p_θ_* : X × C → P(X), which, given an unperturbed cell image *x*_0_ ∼ *p*_0_ and a perturbation *c* ∈ C, predicts the resulting conditional distribution *p*(*x*_1_|*x*_0_*, c*). From this distribution, new images can be sampled to simulate the effects of the perturbation, such that *x*_1_ ∼ *p*_1_ (Fig. 1a).

The input space consists of multi-channel microscopy images, where X ⊂ R*^H^*^×^*^W^* ^×^*^C^*. Here, *H* and *W* represent the image height and width, while *C* denotes the number of channels, each highlighting different cellular components through specific fluorescent markers (analogous to RGB channels in natural images, but capturing biological structures like mitochondria, nuclei, and cellular membranes).

The perturbation space C includes two types of biological interventions: chemical (drugs) and genetic (gene modifications). Chemical perturbations involve compounds that target specific cellular processes — for example, affecting DNA replication or protein synthesis. Genetic perturbations can turn off gene expression (CRISPR) or upregulate gene expression (ORF).

This generative model enables in silico simulation of cellular responses, which traditionally require time-intensive and costly wet-lab experiments. Such computational modeling could revolutionize drug discovery by enabling rapid virtual drug screening and advance personalized medicine through digital cell twins for treatment optimization.

#### Data

Cell morphology data are collected through high-content microscopy screening (Fig. 1b) [30]. In this process, biological samples are prepared in multi-well plates containing hundreds of independent experimental units (wells). Selected wells receive interventions — either chemical compounds or genetic modifications — while control wells remain unperturbed. After a designated period, cells are fixed using chemical fixatives and stained with fluorescent dyes to highlight key structures like the nucleus, cytoskeleton, and mitochondria. An automated microscope then captures multiple images per well. This process is called cell painting. Modern automated high-content screening systems have enabled large-scale data collection, resulting in datasets of terabyte to petabyte images from thousands of perturbation conditions [13, 31].

However, the cell painting process has limitations: cell painting requires cell fixation, which is destructive, making it impossible to observe the same cells dynamically during a perturbation. This creates a fundamental constraint: we cannot obtain paired samples {(*x*_0_*, x*_1_)} showing the exact same cell without and with treatment. Instead, we must work with unpaired data ({*x*_0_}, {*x*_1_}), where {*x*_0_} represents control images and {*x*_1_} represents treated images, to learn the conditional distribution *p*(*x*_1_|*x*_0_*, c*).

One solution is to leverage the distribution transformation from control cells to perturbed cells within the same batch to learn conditional generation. Control cells serve as a crucial reference by providing prior information to separate true perturbation effects from confounding factors such as batch effects. Variations in experimental conditions across different runs (batches) introduce systematic biases unrelated to the perturbation itself. For instance, images from one batch may consistently differ in pixel intensities from those in another. Therefore, meaningful comparisons require analyzing treated and control samples from the same batch. As shown in Fig. 1b, this approach helps distinguish true biological responses, like changes in nuclear size, from batch-specific artifacts, like changes in color.

#### Mathematical Formulation

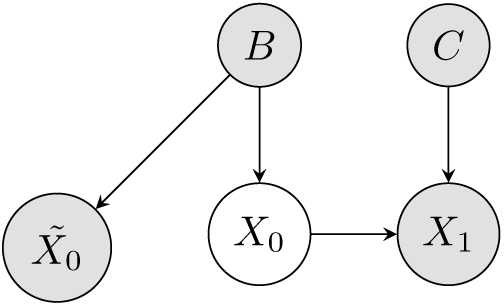

Let us formalize our learning problem in light of the experimental constraints described before. Our objective is to learn a conditional distribution *p*(*x*_1_|*x*_0_*, c*) that models the cellular response to perturbation. However, due to the destructive nature of imaging, we cannot observe paired samples {(*x*_0_*, x*_1_)}. We propose a probabilistic graphical model to address this challenge.

In our graph, random variable *B* denotes the experimental batch, *C* denotes the perturbation condition, *X*_0_ represents the unobservable basal cell state, 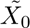 represents control cells from the same batch, and *X*_1_ denotes the perturbed cell state. From our experimental setup, we have access to the control distribution 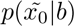 from unperturbed cells and the perturbed distribution *p*(*x*_1_|*c, b*) from treated cells.

We propose to leverage the distributional transition from 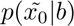 to *p*(*x*_1_|*c, b*) to learn the individual-level trajectory *p*(*x*_1_|*x*_0_*, c*), as shown in Fig. 1c. There are two key reasons. First, there exists a natural connection between *p*(*x*_1_|*c, b*) and *p*(*x*_0_|*b*) through the marginalization *p*(*x*_1_|*c, b*) = *p*(*x*_1_|*x*_0_*, c*)*p*(*x*_0_|*b*)*dx*_0_. Second, while *p*(*x*_0_|*b*) is not directly tractable, we can approximate it using 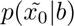 since both the ground-truth *X*_0_ distribution and control distribution 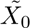 follow the same batch-conditional distribution: *x*_0_ ∼ *p*(·|*b*) and 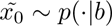.

Our approach of learning 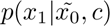 by conditioning on same-batch control images improves upon existing methods that ignore control cells and learn only *p*(*x*_1_|*c*). Intuitively, conditioning on 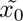 allows the model to initiate the transition from a distribution more closely aligned with the underlying *x*_0_, leading to a better approximation of the true distribution *p*(*x*_1_|*x*_0_*, c*). We formalize this intuition in the following proposition, with proof provided in Appendix:

*Proposition 1* Given random variables *B*, *C*, *X*_0_, 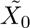, and *X*_1_ following the graphical model above with joint distribution 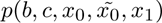, the distribution *p*(*x*_1_|*x*_0_*, c*) can be better approximated by the conditional distribution 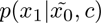 than *p*(*x*_1_|*c*) in expectation. Formally,

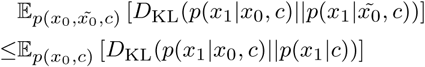

### Enhanced Flow Matching Methodology

As detailed in the previous section, we predict cell morphological changes by transforming distributions between control and perturbed cells under specific conditions within the same batch. In this section, we introduce *CellFluxV2*, which leverages flow matching, a principled framework for learning continuous transformations between probability distributions. We first introduce the basics of flow matching, including how to train and how to perform inference. Then we describe the data sparsity challenge and explain how we address it with three methodology improvements.

#### Flow Matching Basics

Flow matching [32, 33] provides a framework to learn transformations between probability distributions by constructing smooth paths between paired samples (Fig. 1d). It models how a source distribution continuously deforms into a target distribution through time, similar to morphing one shape into another.

More formally, consider probability distributions *p*_0_ and *p*_1_ defined on a metric space (X *, d*). Given pairs of samples from these distributions, flow matching learns a time-dependent velocity field using a neural network *v_θ_* : X × [0, 1] → X that describes the instantaneous direction and magnitude of change at each point. The transformation process follows an ordinary differential equation:

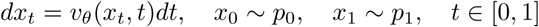

To model perturbation conditions, we extend flow matching by conditioning on perturbations *c* ∈ C. While the source distribution *p*_0_ represents unperturbed cell images, the target distribution now becomes condition-dependent perturbed cell images, denoted as *p*_1_(*x*|*c*). Our goal is to learn a conditional velocity field *v_θ_* : X ×[0, 1]×C → X that captures perturbation-specific transformations [34]:

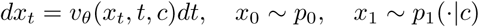

During training, we construct a probability path that connects samples from the source (*p*_0_) and target (*p*_1_) distributions (Fig. 1a). We employ the rectified flow formulation, which yields a simple straight-line path [35]:

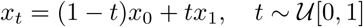

This linear path has a constant conditional velocity field *v*(*x_t_, t, c*) = *dx_t_/dt* = *x*_1_ − *x*_0_, which represents the optimal transport direction at each point. The neural network *v_θ_* is trained to approximate this optimal conditional velocity field by minimizing:

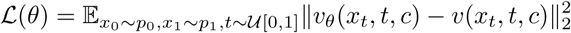

At inference time, given a sample *x*_0_ ∼ *p*_0_ and a perturbation condition *c* ∈ C, we generate *x*_1_ by solving the ODE (Fig. 1b), whose solution is:

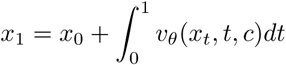

We employ numerical integrators like Euler method or more advanced methods such as Runge-Kutta to solve the ODE.

A well-known method for improving the conditional generation fidelity of flow matching is classifier-free guidance [36], which we incorporate in our framework. During training, we randomly mask conditions with probability *p_c_*, replacing *c* with a null token ∅. At inference time, we interpolate between conditional and unconditional predictions:

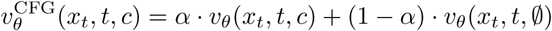

where *α >* 1 controls the guidance strength. We then use 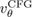 to solve the ODE and obtain *x*_1_.

The velocity field *v_θ_* in *CellFluxV2* is implemented using a Transformer architecture [37]. In contrast to the U-Net design employed in CellFluxV1 [38], Transformers offer improved performance and scalability, enabling us to scale the model to uncover biological scaling laws. Concretely, the Transformer takes latent representations R*^H^*^×^*^W^* ^×^*^C^* as input, where *H*, *W*, and *C* denote the latent height, width, and number of channels. It partitions the input into 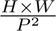 tokens using a patch size *P* and processes these tokens through multiple self-attention layers that capture both local and global image features. The model then outputs a velocity field with the same spatial dimensions R*^H^*^×^*^W^* ^×^*^C^* as the input.

For conditioning, the time variable *t* is encoded using Fourier features, while the condition *c* is first embedded using pretrained representations (described later) and then passed through a learnable linear layer. The time and condition embeddings are combined to form a conditioning signal, which is injected into the Transformer to guide the generation process [34].

#### Addressing Data Sparsity

Flow matching is, in principle, designed to transport an arbitrary source distribution to an arbitrary target distribution. This generality has not been deeply explored because few machine learning problems naturally fit this formulation. Most existing applications use a Gaussian source distribution, which is smooth and unbounded. By framing perturbation modeling as a distribution-to-distribution task, our work is among the first to apply flow matching in a real-world setting where both source and target are empirical distributions.

In our setting, both *p*_0_ (unperturbed control cells) and *p*_1_ (perturbed cells) are empirical and supported by limited observations. Directly applying flow matching performs poorly due to data sparsity. The learned velocity field tends to overfit and fails to generalize in high-dimensional space. To address this, we introduce three methodological improvements: 1) reduce the effective dimensionality of the data; and 2) augment the data and training process to encourage smoother and more generalizable velocity fields. Two of these improvements are unique contributions of this work.

**Latent-Space Modeling.** Reducing dimensionality is a natural way to mitigate data sparsity. We pre-train a variational autoencoder (VAE) that embeds Cell Painting images into a lower-dimensional latent space, and then learn flow matching in that space.

The VAE consists of a standard CNN encoder–decoder. The encoder maps images of size 192 × 192 × 5 (five Cell Painting channels) to a latent representation of size 24 × 24 × 8. The latent dimensionality (8) is a hyperparameter, and we find that 8 works best. The decoder reconstructs the original images from this representation. The VAE is trained with two objectives: 1) a reconstruction loss that provides the main training signal by measuring the discrepancy between the input and reconstructed images; and 2) an adversarial loss, where a discriminator distinguishes real patches from reconstructed patches. Joint adversarial training sharpens high-frequency details and improves overall reconstruction fidelity.

The final reconstruction FID is very low (2.9), which shows that the images can be embedded with roughly a 64× reduction in dimensionality without loss of information. Using this VAE latent space reduces the flow-matching FID from 18.7 to 14.6, a 21.9% improvement, and demonstrates that dimensionality reduction is an effective strategy for overcoming data sparsity.

**Two-Stage Training.** In most machine learning applications of flow matching, only *p*_1_ is an empirical distribution, whereas *p*_0_ is a Gaussian prior that mitigates data sparsity. In contrast, our setting requires learning a transport between two empirical distributions. We introduce a two-stage training scheme that first familiarizes the model with the target distribution before learning the full source-to-target mapping.

The training proceeds in two stages: **Stage 1 (noise** → **target):** Map Gaussian noise to the target images under the same conditioning used in the final task; **Stage 2 (control** → **perturbed):** Map the empirical source distribution to the perturbed target distribution. Both stages use the same flow-matching objective and are trained sequentially for 100 epochs each.

Empirically, Stage 1 acts as a curriculum that stabilizes Stage 2 and smooths the learned velocity field. This two-stage procedure improves sample quality and reduces FID from 14.6 to 10.2, a 30.1% gain. To our knowledge, this training strategy has not been previously explored in flow-matching literature and may motivate further study. **Noisy Augmentations.** Data augmentation provides another way to address data sparsity. We inject random Gaussian noise into both the source images and the interpolation path used by flow matching.

For each source image *x*_0_ ∼ *p*_0_, we apply random noise with probability *p_e_*:

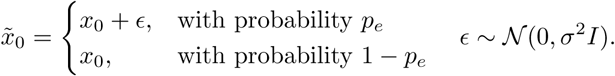

We also replace the deterministic interpolation path

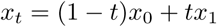

with a noisy interpolant

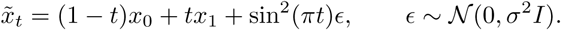

The noise scale *σ* and probability *p_e_* control the local variability that the model observes during training. Intuitively, the injected noise exposes the model to a neighborhood around the interpolation trajectory and discourages overfitting to a single path. This improves robustness and encourages a smoother velocity field in the ambient space. We find that noise augmentation reduces FID from 10.2 to 7.9, a 22.5% improvement, and represents another methodological contribution of this work.

The entire *CellFluxV2* algorithm is summarized in Appendix.

### Data Processing

In this section, we present detailed data processing used by *CellFluxV2*.

#### Datasets used for baseline comparison

For baseline comparison, we use the same datasets processed by Palma et al. [26] to enable fair and head-to-head comparison.

Specifically, these processed datasets include three cell imaging perturbation datasets: BBBC021 (chemical perturbation) [39], RxRx1 (genetic perturbation) [40], and the JUMP dataset (combined perturbation) [13]. The preprocessing protocol involves correcting illumination, cropping images centered on nuclei to a resolution of 96×96, and filtering out low-quality images [26]. The resulting datasets include 98K, 171K, and 424K images with 3, 6, and 5 channels, respectively, from 26, 1,042, and 747 perturbation types. Examples of these images are provided in Fig. 2e.

#### Datasets used for scaling, generalization, and capability study

To study scaling behavior and generalization, we processed a large JUMP-pilot dataset that yielded 11.6 million images after preprocessing. This is roughly 30 times larger than the previously used dataset (the JUMP used for baseline comparison only contains 424K processed images).

For preprocessing, each original JUMP-pilot image first underwent plate-wise illumination correction, followed by channel-wise min–max normalization. We then performed cell-centered segmentation and cropped fixed-size patches of 192×192 pixels. This pipeline produced 11.6 million cell images.

We next constructed the train and evaluation splits. After excluding plates with antibiotics or other anomalies, JUMP-pilot contains 16 chemical perturbation plates, 16 CRISPR plates, and 8 ORF plates. We first split at the plate level, designating two chemical plates and two genetic plates as held-out plates to evaluate out-of-distribution (OOD) generalization across imaging conditions. We then performed a perturbation-level split. JUMP-pilot includes 816 unique chemical and genetic perturbations; we randomly assigned 90 percent to the training set and 10 percent to the test set to evaluate OOD generalization across perturbations. Crossing the plate-level and perturbation-level splits yields four evaluation regimes that enable comprehensive assessment of model capability and generalization: (1) in-distribution (ID) plates with ID perturbations, (2) ID plates with OOD perturbations, (3) OOD plates with ID perturbations, and (4) OOD plates with OOD perturbations. Within the ID plate and ID perturbation regime, wells were further split 90 percent for training and 10 percent for testing for both control and treated wells. The model was trained only on the 90 percent ID plate and ID perturbation data and evaluated on all four regimes. We audited all splits to ensure that no (plate, perturbation) combinations leaked across train and test and that plate and perturbation assignments remained orthogonal.

This large-scale dataset and its structured splits form a robust benchmark for studying model scaling laws and generalization performance in high-content imaging.

### Experimental Details

We describe here the experimental setup, including evaluation metrics, baselines, condition encoding, compute scaling, and hyperparameters.

#### Evaluation metrics

We assess all methods using three categories of metrics. (1) **FID and KID** (lower is better), which quantify distributional similarity between generated and real images using Fréchet and kernel-based distances. Scores are computed on 5,000 generated images for BBBC021 and on 100 randomly selected perturbation classes for RxRx1 and JUMP. We report both overall scores across all samples and class-conditional scores per perturbation. (2) **Mode of Action (MoA) classification accuracy and F1** (higher is better), which measure biological fidelity. A pretrained classifier predicts a compound’s MoA from generated images, and predictions are compared to canonical MoA annotations. (3) **MoA mAP**, computed by extracting CellProfiler features and evaluating retrieval performance. mAP reflects whether images generated under the same MoA rank ahead of images with different MoAs.

#### Baselines

We compare our method to PhenDiff [25] and IMPA [26], the only existing approaches that explicitly incorporate control images into the model design, a setup essential for separating true perturbation-induced changes from nuisance variation such as batch effects. PhenDiff uses a diffusion process that maps control images to noise and then maps noise to target images. IMPA uses a GAN with an AdaIN layer to transfer the style of control images to target images and is tailored to paired image-to-image translation. Our method uses flow matching, which is designed for distribution-to-distribution mapping and is therefore well aligned with the underlying problem. All baselines are reproduced using their official implementations.

#### Condition encoding

Condition representations follow IMPA [26]. Chemical perturbations use concatenated RDKit fingerprints (1024*d*) and Gene2Vec embeddings (200*d*). Genetic perturbations use mean-pooled last-layer embeddings from the ESM2 language model (512*d*) concatenated with Gene2Vec embeddings (200*d*). These standardized embeddings capture domain-relevant chemical or genetic properties.

#### Training details

*CellFluxV2* uses a Transformer-based velocity field trained in two stages. For baseline comparisons, we train for 100 epochs in stage 1 (noise-to-target) and 100 epochs in stage 2 (source-to-target) on 4 A100 GPUs for BBBC021, RxRx1, and JUMP. Each experiment completes within one day.

For scaling experiments, compute grows approximately linearly with model size *N* and dataset size *D*:

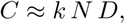

where *k* absorbs implementation and hardware factors. Our largest model (XL, 675M parameters) trained on the full JUMP dataset (11.6M images) for seven days on 4 A100 GPUs. Halving *N* or *D* reduces compute by roughly half.

We perform a careful hyperparameter search on small-scale settings. The best configuration uses AdamW, a linear learning-rate schedule, and a batch size of 128, which we adopt for all other experiments.

**Table 1.**
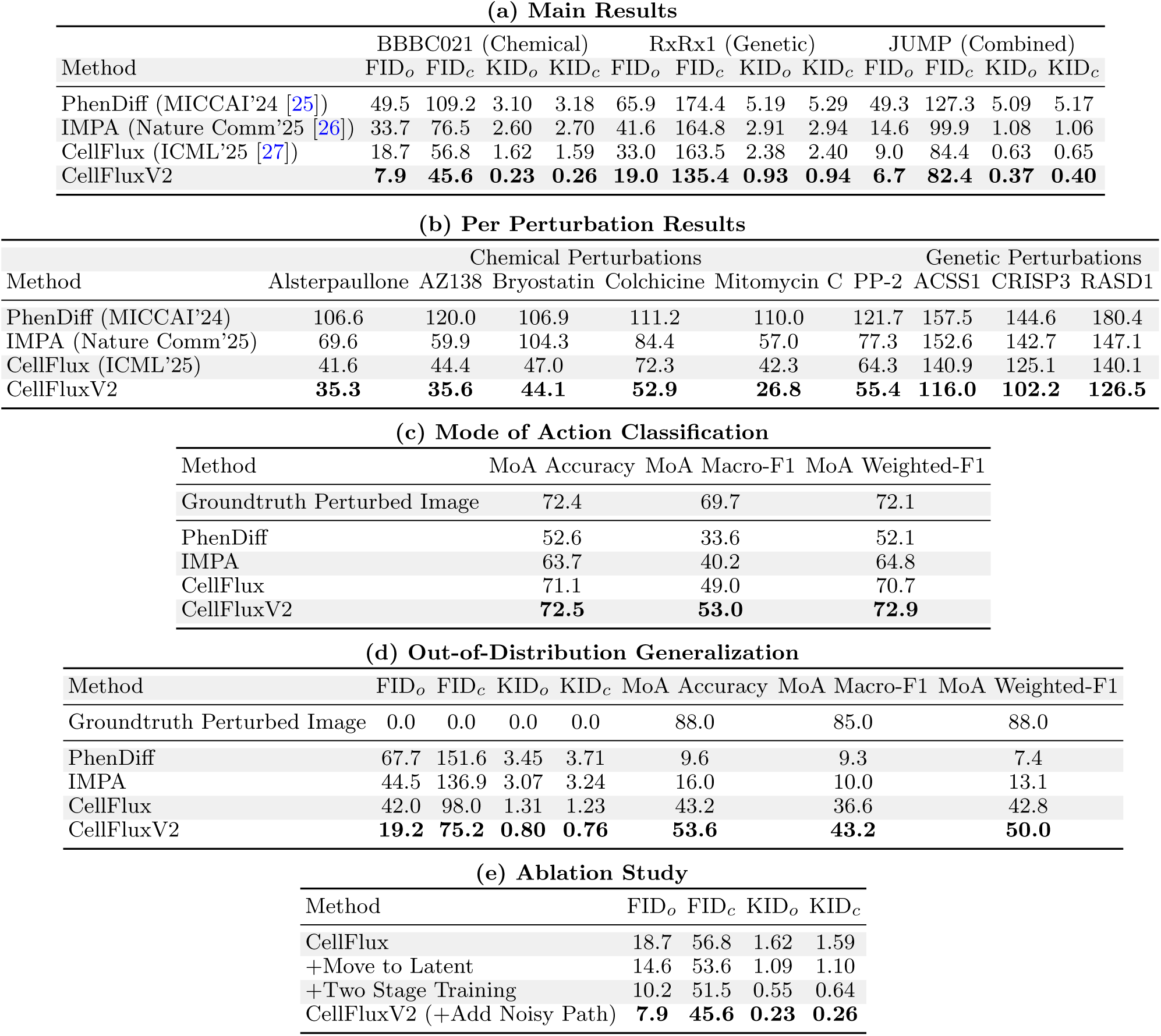
Evaluation of *CellFluxV2*. *(a) Main results. CellFluxV2* outperforms GAN- and diffusion-based baselines, achieving state-of-the-art performance in cellular morphology prediction across three chemical, genetic, and combined perturbations datasets. Metrics measure the distance between generated and ground-truth samples, with lower values indicating better performance. FID*_o_*(overall FID) evaluates all images, while FID*_c_* (conditional FID) averages results per perturbation *c*. KID values are scaled by 100 for visualization. *(b) Per perturbation results.* For six representative chemical perturbations and three genetic perturbations, *CellFluxV2* generates significantly more accurate images that better capture the perturbation effects than other methods, as measured by the FID score. *(c) MoA classification.* On the BBBC021 dataset, we train a classifier to predict the drug’s mode of action (MoA) from cell morphology images and evaluate the accuracy/F1 of generated images. *CellFluxV2* achieves significantly higher accuracy/F1 than other methods, closely aligning with ground-truth images and effectively reflecting the biological effects of perturbations. *(d) Out-of-distribution generalization. CellFluxV2* maintains strong performance when generating cell morphology images for novel chemical compounds not seen during training on BBBC021. *(e) Ablation study.* Each component contributes to *CellFluxV2*’s final performance, emphasizing its importance.

**Table 2.** Biological scaling laws. We establish the first empirical scaling laws in biology, demonstrating that model performance (FID) improves substantially with both increasing data scale (top) and model size (bottom). Beyond in-distribution performance, scaling also enhances the model’s ability to generalize to out-of-distribution image sources and perturbations, highlighting its potential as a robust foundation for virtual cell modeling.

|  | ID Drug, ID Source | ID Drug, OOD Source | OOD Drug, ID Source | OOD Drug, OOD Source |
| --- | --- | --- | --- | --- |
| 3% | 18.31 | 17.37 | 17.82 | 16.72 |
| 10% | 9.55 | 9.41 | 9.95 | 9.52 |
| 30% | 7.10 | 7.03 | 7.60 | 7.53 |
| 50% | 6.70 | 6.65 | 7.28 | 7.35 |
| 100% | 5.98 | 6.00 | 6.38 | 6.86 |
| Tiny | 5.67 | 5.85 | 7.21 | 7.45 |
| Small | 6.29 | 6.34 | 6.88 | 7.22 |
| Base | 5.98 | 6.00 | 6.38 | 6.86 |
| Large | 4.83 | 4.98 | 4.95 | 5.78 |
| X-Large | 4.64 | 4.76 | 4.96 | 5.57 |

## Author contributions

S.Y., E.L., and Y.Z. designed the study. Y.Z., Y.S., T.L., C.W., and L.Z. developed the methodology. Y.Z., Y.S., S.S., and H.L. performed the core model development. Y.Z., Y.S., and Z.W. processed data. Y.Z., Y.S., and Z.W. conducted result analysis. Y.Z., Y.S., Z.W., J.B., A.L., J.N., and D.D. wrote the paper. S.Y. and E.L. jointly supervised the project.

## Data and code availability

All data and code are available at https://github.com/yuhui-zh15/CellFluxV2.

## Acknowledgments

We acknowledge funding from a Stanford Institute for Human-Centered AI Hoffman-Yee Research Grant (S.Y., E.L.) and Param Hansa Philanthropies (E.L.), and funding as well as computing resources from the Chan-Zuckerberg Initiative (E.L.) and Stanford Institute for Human-Centered AI. S.Y. and E.L. are Chan Zuckerberg Biohub — San Francisco Investigators.

## A Theory Proof

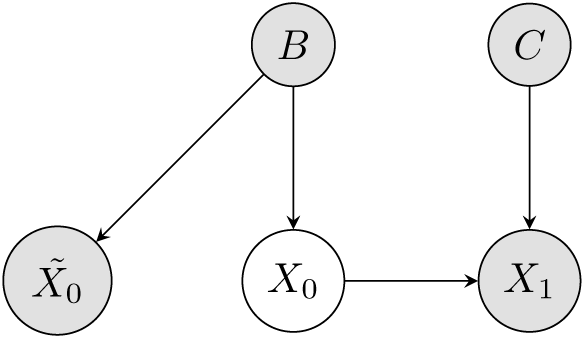

*Proposition 1* Given random variables *B*, *C*, *X*_0_, 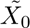, and *X*_1_ following the graphical model above with joint distribution 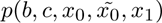, the distribution *p*(*x*_1_|*x*_0_*, c*) can be better approximated by the conditional distribution 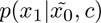 than *p*(*x*_1_|*c*) in expectation. Formally,

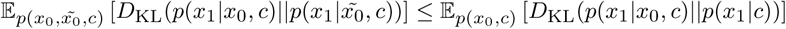

*Proof* According to the definition of conditional mutual information, the term on the right-hand side can be expressed as:

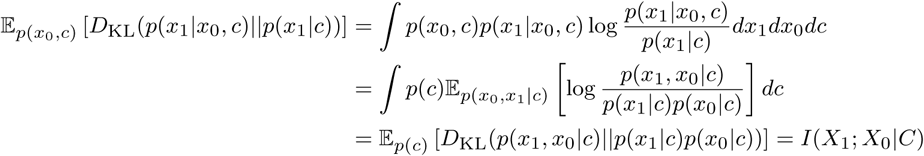

Based on the graphical model, we have the conditional independence 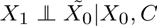*, C*. Thus, we have 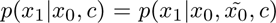. Similarly, we can express the term on the right-hand side as conditional mutual information:

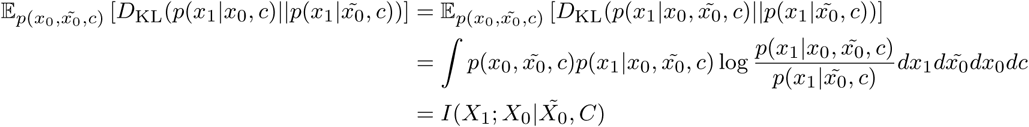

Further, based on the property of conditional mutual information, we have

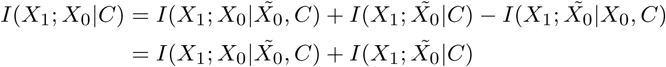

where the second equality is due to the conditional independence relationship 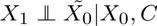, and 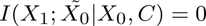

Therefore,

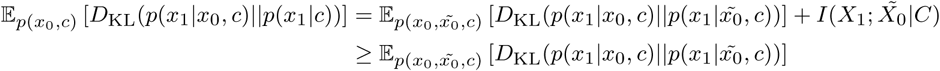

The inequality holds strictly when 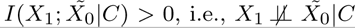, which generally holds true when batch effect exists and variables *X*_0_ and 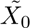 are associated by *B* according to the graphical model.

## B *CellFluxV2* Algorithm

### Algorithm 1

*CellFluxV2* Algorithm

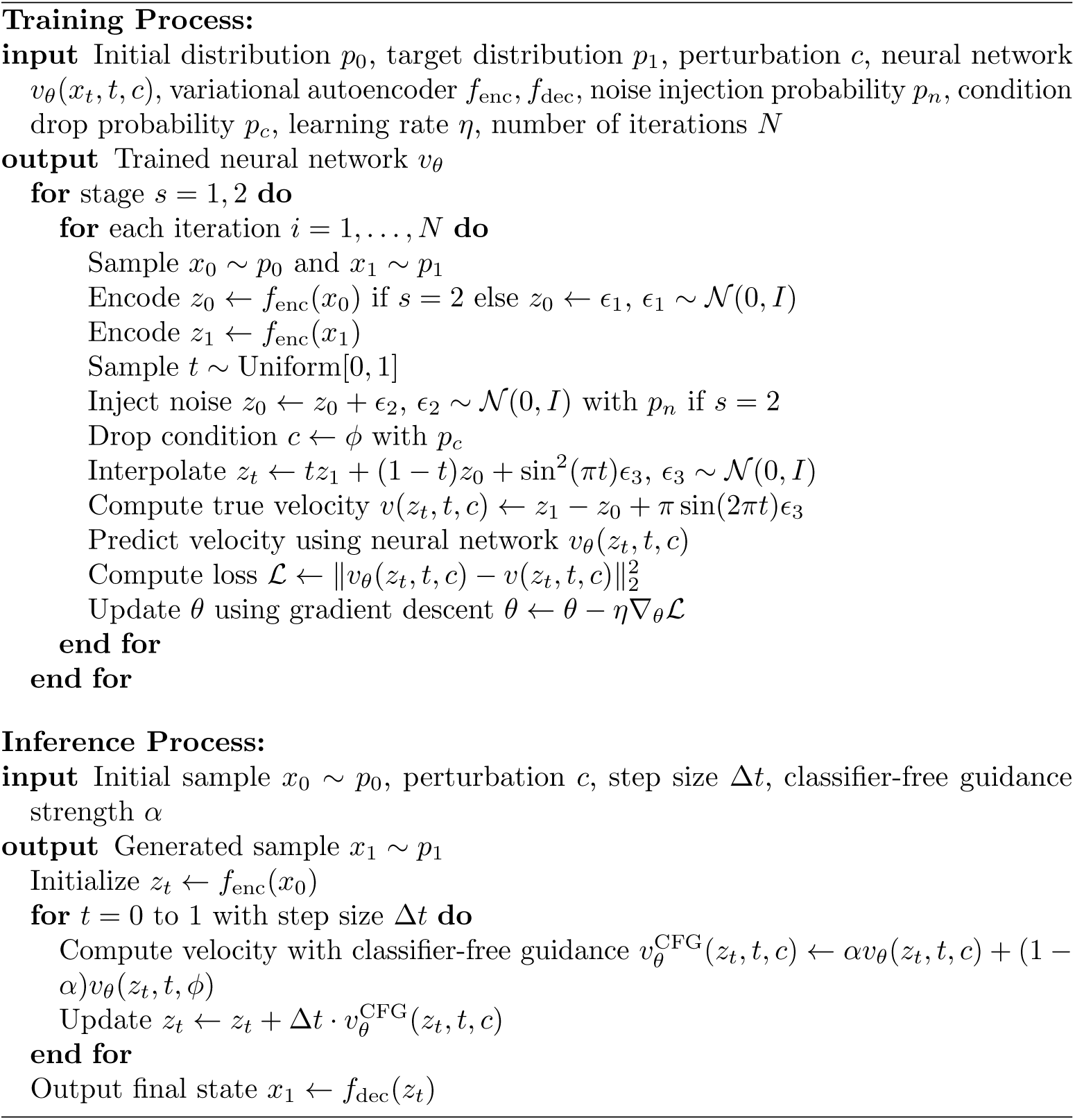

